# Neural ADMIXTURE: rapid population clustering with autoencoders

**DOI:** 10.1101/2021.06.27.450081

**Authors:** Albert Dominguez Mantes, Daniel Mas Montserrat, Carlos D. Bustamante, Xavier Giró-i-Nieto, Alexander G. Ioannidis

## Abstract

Characterizing the genetic substructure of large cohorts has become increasingly important as genetic association and prediction studies are extended to massive, increasingly diverse, biobanks. ADMIXTURE and STRUCTURE are widely used unsupervised clustering algorithms for characterizing such ancestral genetic structure. These methods decompose individual genomes into fractional cluster assignments with each cluster representing a vector of DNA marker frequencies. The assignments, and clusters, provide an interpretable representation for geneticists to describe population substructure at the sample level. However, with the rapidly increasing size of population biobanks and the growing numbers of variants genotyped (or sequenced) per sample, such traditional methods become computationally intractable. Furthermore, multiple runs with different hyperparameters are required to properly depict the population clustering using these traditional methods, increasing the computational burden. This can lead to days of compute. In this work we present Neural ADMIXTURE, a neural network autoencoder that follows the same modeling assumptions as ADMIXTURE, providing similar (or better) clustering, while reducing the compute time by orders of magnitude. Indeed, the equivalent of one month of continuous compute can be reduced to hours. In addition, Neural ADMIXTURE can include multiple outputs, providing the equivalent results as running the original ADMIXTURE algorithm many times with different numbers of clusters. Our models can also be stored, allowing later cluster assignment to be performed with a linear computational time. The software implementation of Neural ADMIXTURE can be found at https://github.com/ai-sandbox/neural-admixture.

## Introduction

The rapid growth in numbers of sequenced human genomes and the proliferation of population-scale biobanks have enabled the creation of increasingly accurate models to predict traits and disease risk based on an individual’s genome. However, different predictive models can be required depending on an individual’s genetic ancestry, and this necessitates accurately characterizing an individual’s genetic ancestry composition at the individual level [1]. Such characterization is also an essential part of most modern population genetic studies and national biobanking projects. However, many existing algorithms for population genetic analyses struggle to keep up with next generation sequencing data sets, where both the number of samples and the number of sequenced positions along the genome, are much greater. This has created an intense need for more computationally efficient and accessible methods for detailed large-scale structure analyses. To date the vast majority of association studies rely on samples from individuals of European descent, thus excluding most of the world’s population and creating a new divide in healthcare [2]. These are the cases of genome-wide association studies (GWAS), which look for correlations between genomic sequences and phenotypes, and predictive models like polygenic risk scores (PRS), which indicate genetic predisposition to phenotypes. The inclusion of fast, interpretable algorithms that characterize the ancestry makeup of genetic sequences is an important part of facilitating the creation of diverse association studies and expanding the reach of personalized genomic medicine.

A common approach for resolving the population structure within a genetic dataset is to describe each sample by a set of proportional cluster assignments obtained through an unsupervised clustering algorithm. Such methods take as input each individual’s sequence of single nucleotide polymorphisms (SNPs), that is, those positions along the genome known to vary between individuals. There are over 10 million known SNPs in the human genome with most of the remainder of the human DNA sequence shared in common between all humans. Such positions are commonly encoded with a binary value, where 0 is used to encode the most common (or reference) variant at that SNP position on the genome, and 1 is used to encode the minority (or sometimes called “alternative”) variant. This binary encoding works, because the vast majority of such variable positions have only one alternative to the common variant (are biallelic). The frequency distribution of these variants, and the correlations (linkage disequilibrium) between neighboring SNPs, will vary between populations due to different founder events, migration histories, and genetic drift experienced by those different populations. These differences can lead to predictive models trained on one population failing when faced with sequences from an unseen population. In addition, characterizing differing variant frequencies between populations can provide valuable historical and demographic information such as divergence times, migration events, and historical census size [3].

Analysis and studies performed in biobank typically involve large amounts of data. If the computation resources permit it, the dataset is processed in its totality at once, otherwise it can be split into multiple parts (commonly still of significantly large size). Depending on the nature of the data, several preprocessing steps can be performed including data filtering and possibly imputation and phasing. After that, methods that capture the population structure of the data, such as ADMIXTURE, are applied to the data. Commonly PCA is also used. The ADMIXTURE coefficients are then used in downstream steps, such as GWAS for which individuals of various ancestral backgrounds may be selected. It is quite common that each processing step needs to be iterated multiple times during the study whenever the genomic sequence data is preprocessed while troubleshooting errors in merging of datasets and refining QC filters. This can be a problem if some of the steps (e.g. ADMIXTURE) are highly computationally consuming. In some scenarios, (*e.g*. tracking of Covid-19 community spread) where new patient genomic sequences become available in a streaming fashion, having inference-only pretrained methods (such as our proposed method), highly speeds the analysis, and removes the need of dealing with large amounts of data (training datasets) every time inference is required.

In this paper we present an autoencoder that implements one of the most widely used clustering methods for population genetics applications: ADMIXTURE [4, 5]. ADMIXTURE was developed as a more computationally efficient solution than STRUCTURE [6], and we now take this pursuit of efficiency to the next generation. Our proposed method, *Neural ADMIXTURE*, follows the same modeling assumptions as ADMIXTURE but re-frames the task as a neural network-based autoencoder, providing much faster computational times, both on GPU and CPU, and higher quality cluster assignments. Additionally, we introduce *Multi-head Neural ADMIXTURE*, which combines multiple decoders to obtain clustering results equivalent to running the original ADMIXTURE repeatedly with different priors for numbers of clusters. Both methods also include a supervised version that performs regular classification given ground truth training labels. The proposed method is fully compatible with the original ADMIXTURE framework, allowing use of ADMIXTURE results as initialization for Neural ADMIXTURE parameters, and vice-versa.

## Related work

Model-based clustering methods such as FRAPPE [7], STRUCTURE [6] and ADMIXTURE [4, 5] are the most commonly used unsupervised clustering techniques for analyzing the population structure of genomic sequences. These methods decompose each input sequence into a set of cluster assignments and compute one explicit centroid for each cluster. This resembles probabilistic versions of non-negative matrix factorizations (NMF). Specifically, the cluster assignments specify what proportion of each ancestry cluster an individual has, while the centroids indicate the SNP variant frequencies at each genetic position for each cluster. These methods allow the user to visualize ancestry composition within genetic datasets, compute how genetically distant different population groups are, and compute statistics that allow the dating of migration history. STRUCTURE [6] makes use of Bayesian models, using a Dirichlet prior for cluster assignments and the cluster centers, trained with Markov chain Monte Carlo (MCMC), making it highly computationally intensive. FRAPPE [7] and ADMIXTURE [4, 5] make use of maximum-likelihood point estimates, obtaining predictions with a quality compared to STRUCTURE, but with much faster computational times. FRAPPE makes use of the Expectation-Maximization algorithm (EM) while ADMIXTURE, as explained in depth in the following section, makes use of a faster block relaxation quasi-Newton optimization technique. However, each method still requires many hours of compute time and is not well suited for modern biobank datasets with tens or hundreds of thousands of samples and millions of SNP feature dimensions.

Several autoencoder architectures similar to our work have been previously proposed. Some examples include the Dirichlet Variational Autoencoder [8], Deep Archetypal Analysis (DeepAA) [9], and Genotype Convolutional Autoencoder (GCAE) [10]. Such networks encode each sample as a point within a convex hull, or as a set of proportions and probabilities. The Dirichlet VAE replaces the commonly used Gaussian prior in the bottleneck by a Dirichlet prior. DeepAA adds constraints to enforce that the bottleneck representation is non-negative and sums to one. GCAE is a convolutional neural network with a Softmax activation in the bottleneck that provides similar clustering results as ADMIXTURE, while being more computationally intensive. Such methods are composed by non-linear encoders and decoders, which deny them the interpretability that our method provides. In fact, our proposed method can be seen as a non-variational version of the Dirichlet VAE with a linear decoder (without bias) and additional constraints in the dynamic range of its weights. In [11], a VAE with a linear reconstruction function (decoder) is explored. While this approach to retain interpretability resembles ours, our work differs by using a deterministic non-generative model. In addition we introduce a multi-head method that performs several cluster assignments (for different numbers of clusters) in only a single forward pass, and implement a supervised approach, among other innovations.

Neural Network-based supervised methods for ancestry classification have also been introduced in the past. Some examples include LAI-Net [12], which provides a high-resolution ancestry estimate along a chromosome sequence, Diet Networks [13], which proposes a genome classifier with different regularization techniques to deal with the high dimensionality of genomic data, and Locator [14], which treats ancestry inference as a geographical prediction problem. While these methods can accurately classify genomes once trained, the ground truth labels used to train these supervised methods are typically hand-crafted reflections of concepts such as ethnicity, or self-reported race of the individual samples. These human-informed classes do not always reflect the full spectrum, or significant clusters, of genetically relevant substructure within and between populations. Therefore, in many genetic applications, it is preferred to use unsupervised methods that do not rely on the complexity of socially-constructed labeling schemes.

## Background

### ADMIXTURE

In this work, we follow the notation presented in [5]. Note that each individual human has two copies of each chromosome (one paternal and one maternal). Therefore, for a given individual at each genomic position we have the possibility of four different combinations of biallelic SNPs (0/0, 0/1, 1/0, 1/1). It is common practice to sum both maternal and paternal sequences, obtaining a count sequence *n_ij_*. In this scenario, an individual *i* has *n_ij_* ∈ {0, 1, 2} copies of the minority SNP *j*. ADMIXTURE models each individual’s sequence, given a fixed number of clusters (population groups) *K*, as *n_ij_* ~ Bin(2, *p_ij_*), where *p_ij_* = ∑*_k_q_ik_f_kj_*, with *q_ik_* denoting the fraction of population *k* assigned to *i*, and *f_kj_* denoting the frequency of SNPs with value “1” *j* in population k. ADMIXTURE applies block relaxation to try to find the parameters *Q* and *F* that minimize the following negative log-likelihood function:

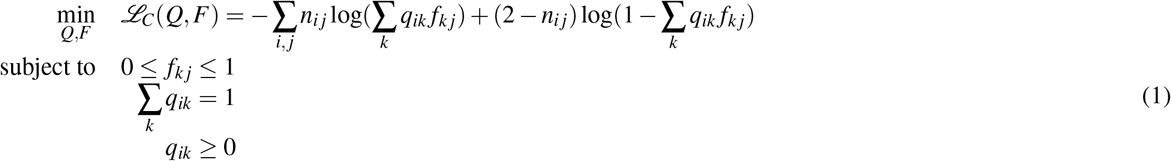

where *Q* = (*q_ik_*) and *F* = (*f_kj_*). ADMIXTURE also allows an expectation-maximization (EM) based optimization, identical to FRAPPE [7], but this approach is slower than the block relaxation approach [4]. The value of *K* is typically chosen by using a cross-validation procedure [5], necessitating runs across a range of values.

ADMIXTURE allows, among others, two valuable optimization alternatives, which are the *projective* analysis and the *supervised* training. In the projective analysis, *F* is initialized to a previously estimated matrix, and only *Q* is optimized. The initialization of *F* may come from previously fit ADMIXTURE models for which the learnt population structure is considered robust. This is especially useful in scenarios where ADMIXTURE is fit in a large dataset and new unseen samples need to be processed. The projective analysis allows estimation of the cluster assignments without the need of fitting the complete model with all the dataset samples. On the other hand, the supervised version requires that some population ancestries are known, so some rows of *Q* are initialized and fixed to these ancestries, while the rest of the rows of *Q* and *F* are optimized normally.

The block relaxation optimization in ADMIXTURE runs much faster than its main competitors, namely FRAPPE [7] and STRUCTURE [6]. Moreover, it can be run in multi-threading mode, greatly boosting the execution time. However, this boost is still insufficient when dealing with either a large number of samples or a large number of SNPs. Our neural network version of the algorithm, however, benefits from massive speedups during training (*e.g*. minibatch training, GPU usage), as well as during inference time with a well-chosen architecture.

## Neural ADMIXTURE

### Network architecture

ADMIXTURE can be formulated as a non-negative matrix factorization problem. Let *X* denote the training samples, where the features are the alternate allele counts per SNP. Then, *X* ≈ *QF*, where *Q* are the assignments, *F* are the alternate allele frequencies per SNP and population, and the negative log-likelihood in Equation (1) is the distance metric between *X* and *QF*. This can be naturally translated into the neural network world as a vanilla autoencoder, with *Q* = Ψ_*θ*_ (*X*) being the bottleneck estimated by the encoder function Ψ (parametrized by a set of weights *θ*) and *F* being the decoder weights themselves. The encoder-decoder architecture is depicted and described in Figure 1. The fact that *Q* is estimated at every forward pass and not learnt as a whole for the training data means that, at inference time, we will not have to run the optimization process again, as in ADMIXTURE’s projective analysis, but instead perform a simple forward pass.

**Figure 1.**
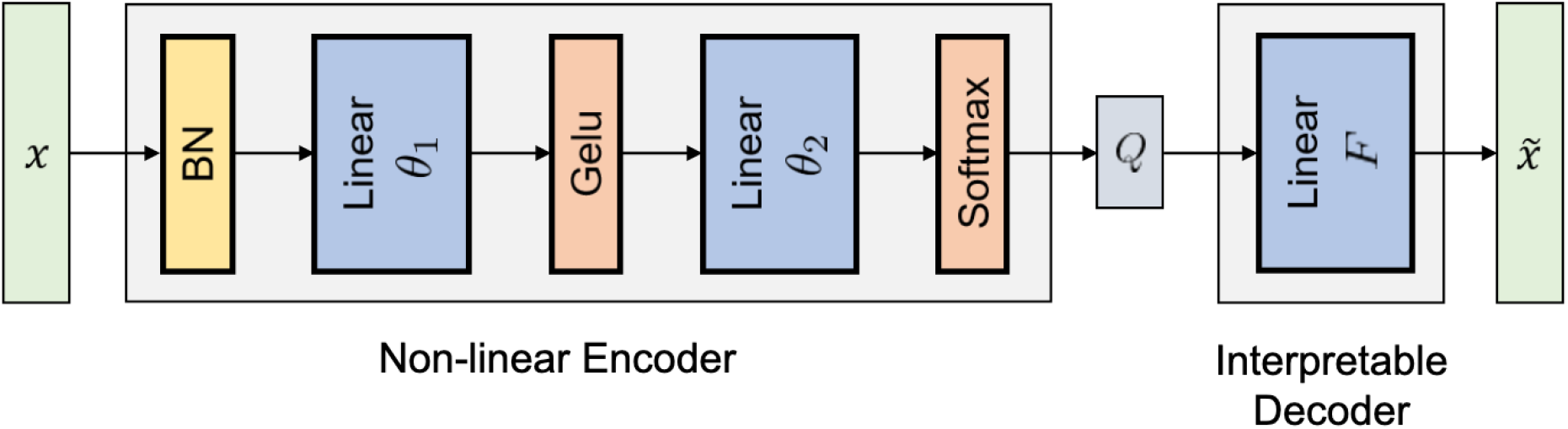
Neural ADMIXTURE architecture. The batch-normalized input sequence is projected into 64 dimensions and processed by a GELU non-linearity. The cluster assignment estimates *Q* are computed by feeding the 64-dimensional sequence to a K-neuron layer activated with a Softmax. Finally, the decoder reconstructs the input using a linear layer with weights *F*. Note that the decoder is restricted to have this architecture to ensure interpretability.

Note that the restrictions in the optimization problem (Equation (1)) impose restrictions in the architecture. Those relating to *Q* (∑*_k_ q_ik_* = 1 and *q_ik_* ≥ 0) can be enforced by applying a softmax activation at the encoder output, making the bottleneck equivalent to the population estimates. Furthermore, while the decoder restriction (0 ≤ *f_kj_* ≤ 1) could also be enforced in the architecture itself (e.g. applying the sigmoid function to the decoder weights), we have found that it suffices to simply project the weights of the decoder to the interval [0, 1] after every optimization step, which is one of the most common forms of projected gradient descent [15].

We note that, critically, the decoder must be linear and cannot be followed by a non-linearity, as it would break the interpretability of the *F* matrix and the equivalence between the decoder weights and the cluster centroids (frequencies per SNP and per cluster) would be lost. On the other hand, the encoder architecture is free from constraints, and it may be composed of several neural layers with its corresponding non-linearities, if deemed appropriate. In fact, the proposed Neural ADMIXTURE includes a 64-dimensional non-linear intermediate layer with a Gaussian Error Linear Unit activation [16] before the bottleneck, as well as a batch normalization layer that acts directly on the input.

The batch normalization operation re-scales the data to have zero mean and unit variance. As the mean for each SNP is its frequency *p* and its standard deviation σ is 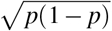, the {0| 1} input gets encoded as 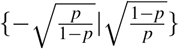, therefore providing more explicitly the information of the allele frequencies to the network.

The ADMIXTURE model does not exactly reconstruct the input data as a regular autoencoder would do, as the input SNP genotype sequences, *n_ij_* ∈ {0, 1, 2}, and the reconstructions *p_ij_* ∈ [0, 1], do not have matching ranges. This can easily be remedied by dividing the genotype counts by two, so that now the input data are 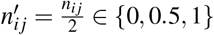. Moreover, instead of minimizing 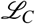 (Equation (1)), we propose minimizing the binary cross-entropy instead, using a penalty term on the Frobenius norm of the first non-linear layer weights, *θ*_1_:

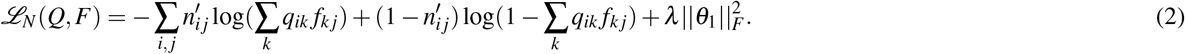

This regularization term avoids hard assignments in the bottleneck, which helps during the training process and reduces overfitting. In Equation (3) we show that the proposed optimization problem and the ADMIXTURE one are equivalent (excluding the regularization term) by using Equations (1) and (2):

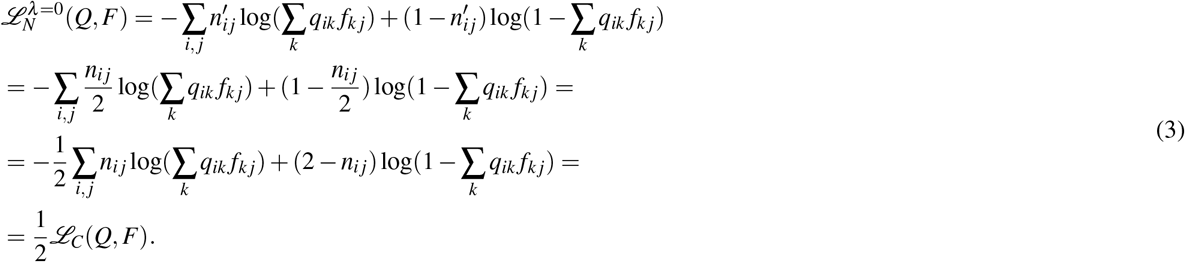

Note that a perfect reconstruction can be obtained by setting the number of clusters (*K*) equal to the number of training samples or to the number of input dimensions. However, we want the bottleneck to capture elementary information about the population structure of the given sequences, therefore we make use of low-dimensional bottlenecks.

### Decoder initialization

Due to the restriction that non-linearities cannot be used in the decoder, as well as the fairly large number of parameters for a single layer, the decoder weights (and thus, the overall performance of the model) are quite sensitive to the initialization. Common initializations, such as Xavier [17], do not work successfully in this architecture. However, the fact that the decoder is interpretable can be exploited in our favour, as we can try to insert information about the population structure into the initialization in an unsupervised manner. As the entries of (*f_kj_* are the frequencies of the alternate variant of SNP *j* in population *k, f_k_* almost coincides with the centroid of the samples in population *k*. This suggests that classical clustering methods can be performed with the results used to initialize the decoder weights.

In the high dimensional space that we work, even fast clustering algorithms such as K-Means would yield high execution times. Instead of clustering in the original feature space, we propose to project the data using Principal Components Analysis (PCA) into a lower dimensional subspace of only a few (2 to 8) principal components and then perform K-Means. PCA is widely used in genetic analyses, as a small number of principal components often explain much of the population substructure of the sequences [18, 19, 20], which is what we are interested in. Hence, to initialize the decoder weights, we propose Algorithm 1:

#### Algorithm 1: PCK-Means Initialization

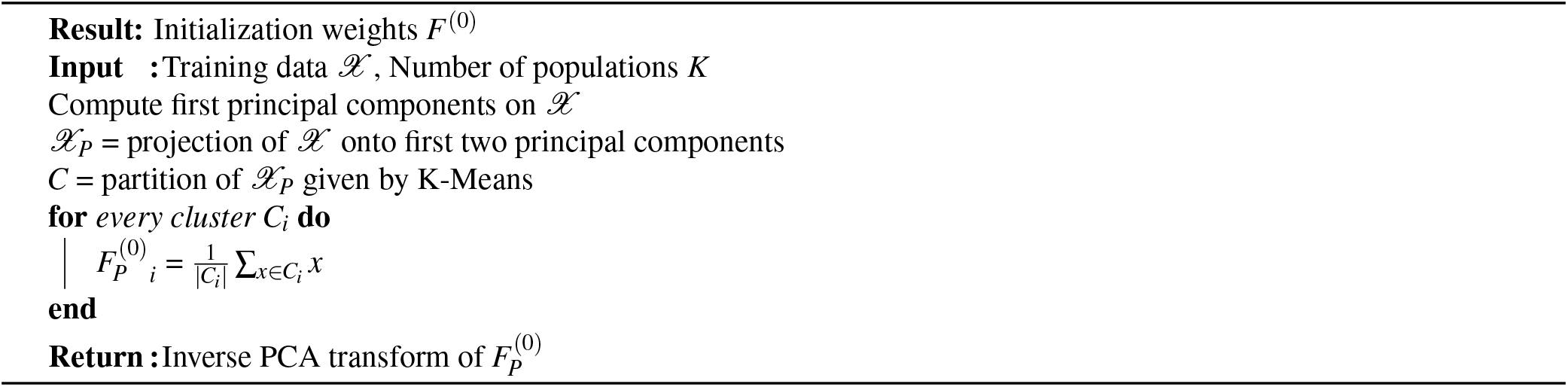

Nevertheless, a large presence of admixed individuals (samples with more than one ancestry linked to them) in the data can lead to degraded performance if Algorithm 1 is used, as *e.g*. the admixed individuals may be identified as a cluster itself depending on the degree of admixture present. Admixed individuals can be considered to be convex combinations of single ancestry individuals, who will typically represents extreme points in a convex polygon. Therefore, a valid approach might be to use archetypal analysis [21] on the projected data. Archetypal analysis can be used to rapidly find representative extreme points (archetypes) in an unsupervised manner, and has already been applied successfully in the context of population genetics [22]. This results in Algorithm 2:

#### Algorithm 2: PCArchetypal Initialization

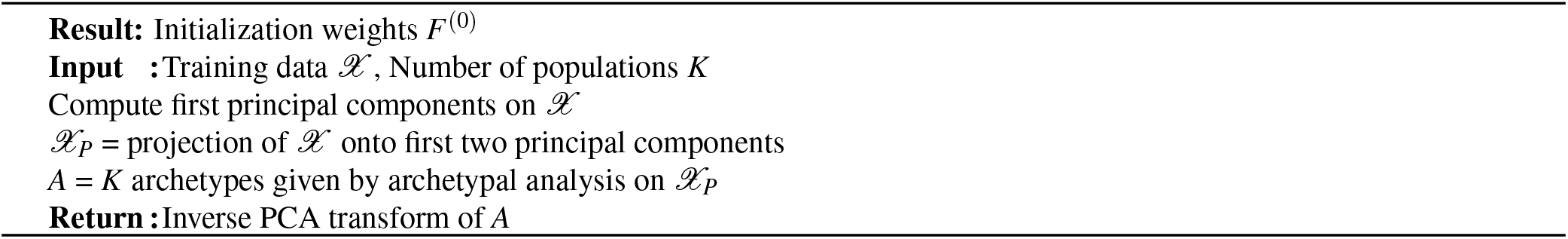

### Multi-head architecture

In ADMIXTURE, cross-validation must be performed in order to choose the number of population clusters (*K*), unless specific prior information about the number of population ancestries is known. Furthermore, in many applications, practitioners desire to observe how cluster assignments change as the number of clusters increase. With the number of both sequenced individuals and variants increasing, the feasible number of different trials of cross-validation rapidly decreases due to its computational cost. As a solution, we propose a variation to Neural ADMIXTURE: the *Multi-head Neural ADMIXTURE* (MNA), which takes advantage of the 64-dimensional latent representation (from now on, shared bottleneck) computed by the encoder. In MNA, the shared bottleneck is jointly learnt for different values of *K*, {*K*_1_ … *K_H_*}.

Figure 2 shows how the shared bottleneck of the multi-head structure is split into *H* different heads. The i-th head consists of a non-linear projection to a *K_i_*-dimensional vector, which corresponds to an assignment assuming there are *K_i_* different populations in the data. While every head could be concatenated and fed through a decoder, this would cause the decoder weights *F* not to be interpretable. Therefore, every head needs to have its own decoder and, thus, *H* different reconstructions of the input are performed in every forward pass.

**Figure 2.**
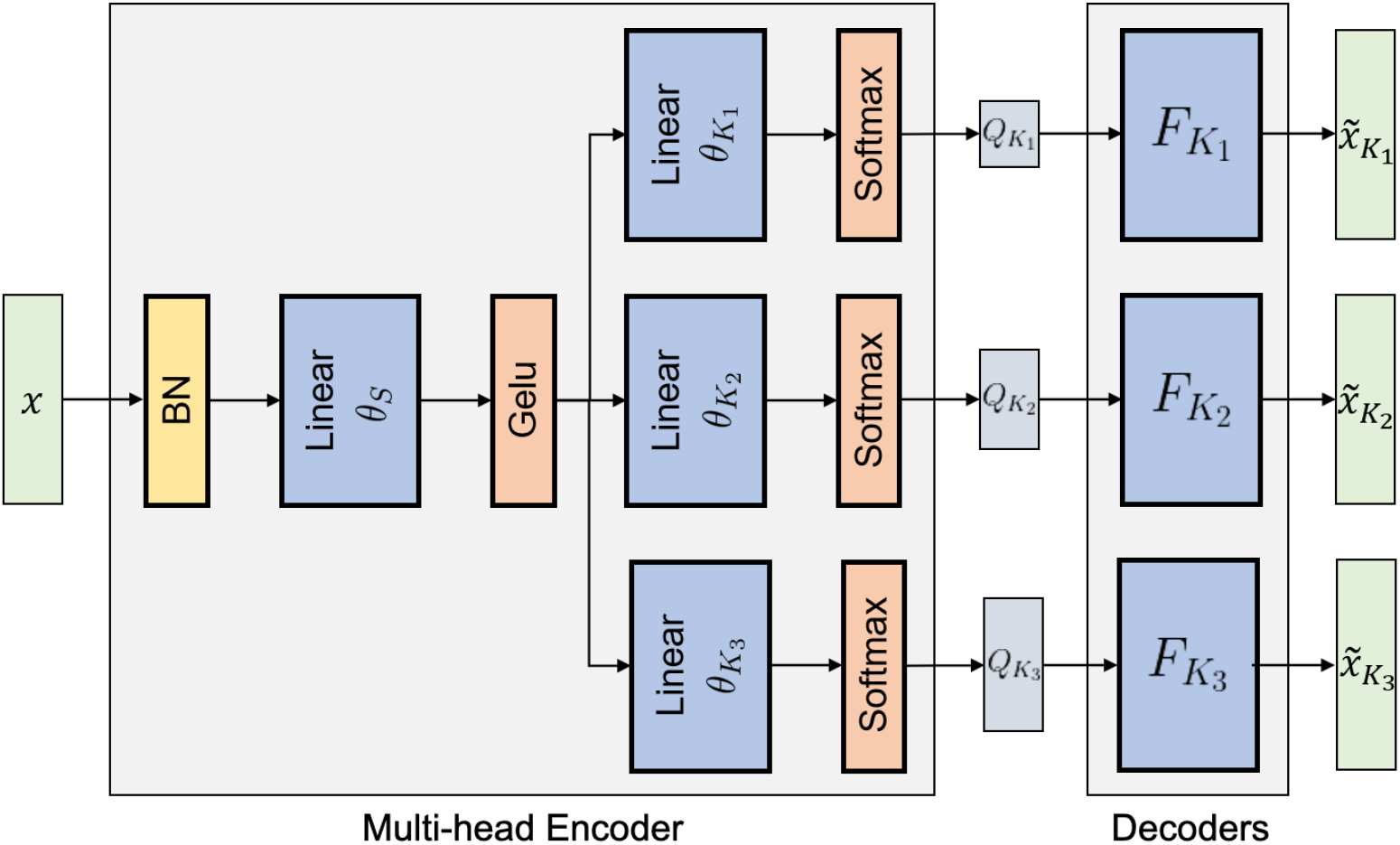
Multi-head Neural ADMIXTURE architecture (*H* = 3).

As we have *H* reconstructions, we will now have *H* different loss values. We can train this architecture by minimizing,

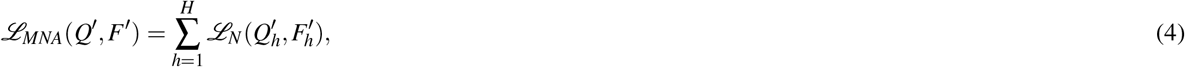

where 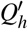 and 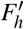 are, respectively, the cluster assignments and the SNP frequencies per population for the *h*-th head. The restrictions of the ADMIXTURE optimization problem (Equation (1)) must be satisfied by 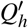 and 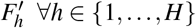.

The multi-head architecture allows computation of *H* different cluster assignments, for different values of *K*, efficiently in a single forward pass. Results can then be both quantitatively and qualitatively analysed in order to decide which value of *K* is the most suitable for the data.

### Supervised training

ADMIXTURE allows for supervised training by fixing some (or all) entries in the *Q* matrix. The same approach cannot be applied to the neural network architecture because *Q* are not learnt parameters but are instead the output of the encoder. As a solution, we propose to add a classification loss row-wise to the *Q* matrix, which correspond to the predicted cluster assignments of every individual. Let *Q_GT_* denote the ground truth ancestries and 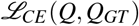 denote the cross-entropy loss. In the supervised version, the optimization problem (assuming a single-head architecture) is formulated as

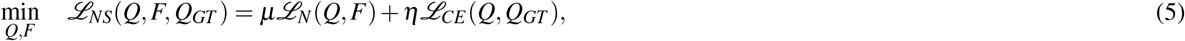

along with the restrictions from Equation (1). Note that unsupervised Neural ADMIXTURE can be seen as a particular case by setting *η* = 0. Furthermore, having both losses allows for semi-supervised training, where only part of the training samples have ground truth labels.

In the supervised learning setting, instead of initializing the decoder weights using PCK-Means (Algorithm 1) or PCArchetypal (Algorithm 2), we can exploit the fact that we know to which population each sample belongs; the decoder weights can simply be initialized as the centroid of each ground truth population, a straightforward computation. Moreover, this initialization will avoid permutation issues (*i.e*. cluster “i” found by the initialization algorithm may not correspond to population encoded as “i”) which would make convergence slower, or even have a negative impact on performance.

### Pretrained Neural ADMIXTURE

As a final contribution, we propose a training scheme which allows reusing the results of a previously optimized interpretable global ancestry method (such as ADMIXTURE) to speedup inference on a novel dataset. This is especially useful when several runs have already been performed, so that full retraining, which is computationally expensive with large datasets, is not desirable.

Let *F_A_* denote the allele frequencies matrix estimated by an arbitrary algorithm. This matrix can be used in Neural ADMIXTURE so the *Q* estimates will be similar to those given by the already trained method by initializing the decoder weights *F*^(0)^ to *F_A_*, and then freezing *F*^(0)^ and learning *Q* in a few epochs.

While the resulting *Q* estimates will not be exactly equal to the estimates coming from the method used to initialize the decoder weights, the computation of cluster assignments will be sped up noticeably.

## Experiments

### Datasets

We use a comprehensive set of publicly available human whole genome sequences from diverse populations across the world, combining the 1000 Genomes Project [23], the Simons Genome Diversity Project [24], and the Human Genome Diversity Project [25], as well as realistic simulated data.

On the one hand, we have compiled single-ancestry data including 550 Africans (AFR), 75 Native Americans (AMR), 651 East Asians (EAS), 496 Europeans (EUR), 27 Oceanians (OCE), 590 South Asians (SAS), and 127 West Asians (WAS). Each category is defined geographically with the American populations additionally filtered to exclude post-colonial groups with recent origins from other continents (e.g. Europe and Africa) by considering only samples with over 95% indigenous local ancestry segments. The genome sequences are from anonymous individuals sequenced with their full consent. These samples are randomly split into train and validation using a 80/20 split. The validation set will be used to compare performances of ADMIXTURE and Neural ADMIXTURE on unseen data. We make use of two different datasets: *Chm-22* and *Chm-22-Sim*. Chm-22 includes the subset of the genome sequence encoded on chromosome 22. Chm-22-Sim is an augmented version of the Chm-22 data: it contains simulated descendants of the real individuals, created using a recombination simulation program, PyAdmix [26] with the simulations performed independently on the train and validation partitions of Chm-22. A total of 400 individuals per ancestry are generated in the training set and 50 in the validation set. Both Chm-22 and Chm-22-Sim have 317,408 SNPs.

We also report results on a single-ancestry dataset (*All-Chms*) consisting of variants from all human autosomes which have been filtered (SNPs with a minor allele frequency (MAF) < 0.05 have been removed) and pruned to minimize linkage disequilibrium (variants which have an *r*^2^ > 0.01 with any other variant in a 50 SNP sliding window), in order to have a dataset which satisfies the assumptions of the models.

On the other hand, to assess performance when there is presence of admixed individuals in a controlled setting, we have chosen single-ancestry individuals for 5 super-populations (AFR, AMR, EAS, EUR, SAS) and generated admixed samples. Moreover, single-ancestry founders were also simulated in order to have balance in the number of classes. This procedure results in the Pruned-Admixed-Balanced (PAB) dataset.

Finally, as commonly reported in related work, we have used a fully synthetic dataset which we have generated using the Pritchard-Stephens-Donnely (PSD) model [6] using the software provided by [27].

### Benchmark setup

In order to have a better review of the performance of Neural ADMIXTURE, we compare the time and quality performance not only against ADMIXTURE, but also against AlStructure [28] and TeraStructure [29], which were designed to improve the scalability of admixture models. TeraStructure iteratively computes *Q* and *F* while avoiding a high computational load by subsampling SNPs at every iteration, which makes the algorithm fast. AlStructure first estimates a low-dimensional linear subspace of the admixture components and then search for a model in the latter subspace which satisfies the modelling constraints, which is a fast alternative to the iterative or maximum likelihood schemes followed by most algorithms.

Moreover, to have a reference against another neural network-based model, we also compare against HaploNet [30]. HaploNet consists of a variational autoencoder (VAE) to map parts of the sequence (windows) to a low-dimensional latent space, on which clustering is then performed using Gaussian mixture priors. While the global structure of the data is preserved in the low-dimensional space, direct interpretability of the allele frequencies is not preserved.

All models are optimized using 16 threads on an AMD EPYC 7742 (x86_64) processor, with 64 cores and 512GB of RAM. To assess GPU performance of Neural ADMIXTURE, all networks are trained on a NVIDIA Tesla V100 SXM2 of 32GB. The same type of graphics card is used to run inference on the trained models.

All algorithms run until the default convergence parameter is achieved. In the case of Neural ADMIXTURE, we have defined that convergence is achieved when the difference in the loss value between two subsequent iterations is less than 10^−^6. The hyperparameters are discussed in the Appendix. The networks are implemented using the PyTorch framework [31].

In the following paragraphs, inference is understood as running a projective analysis on the validation set in ADMIXTURE, and performing a forward pass to obtain the *Q* estimates in Neural ADMIXTURE. Let *N* denote the number of samples and *M* the number of variants (SNPs). In order to assess performance of the *Q* estimates, we match the assignments with the ground truth assignments and report the root mean squared error (RMSE) between them

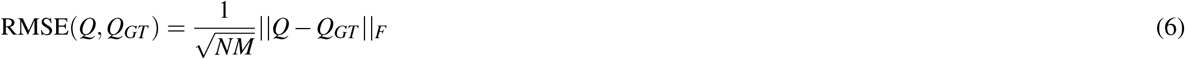

as well as the RMSE between the ground truth allele frequencies (*F_GT_*) and the estimated frequencies (*F*)

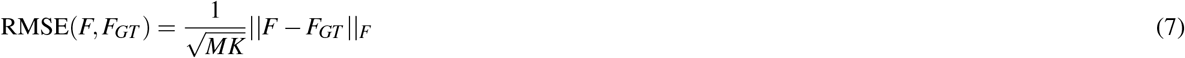

. Furthermore, we propose a new metric, Δ, defined as

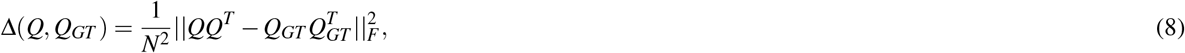

which is equivalent to computing the mean squared difference between the covariance matrices of both the estimated and the target population. In case the estimates *Q* completely agree with *Q_GT_* (up to permutation), then Δ will be 0. The more the disagreement, the higher the value of Δ.

We are interested in these metrics as they are more easily interpretable than the loss function value itself. We are aware that these pseudo-supervised metrics will not give us the true quality of the predictions of the models, as some labels may not be accurate. However, assuming that most of the labels are correct, it will allow us to quantitatively analyze the agreement between the handcrafted labels and the model estimates, and therefore give us an estimation of the quality of the predictions, which we will use to compare among the different algorithms.

### Single-head results

#### Training data

The results in Table 2 and Figure 3 show how Neural ADMIXTURE is systematically faster than the rest of the algorithms on both CPU and GPUs. Taking into account the fact that typically several runs with different values of *K* must be run, the speedup would become even higher by exploiting the Multi-head Neural ADMIXTURE architecture. Some results using this architecture are described in the following sections.

**Table 1.**
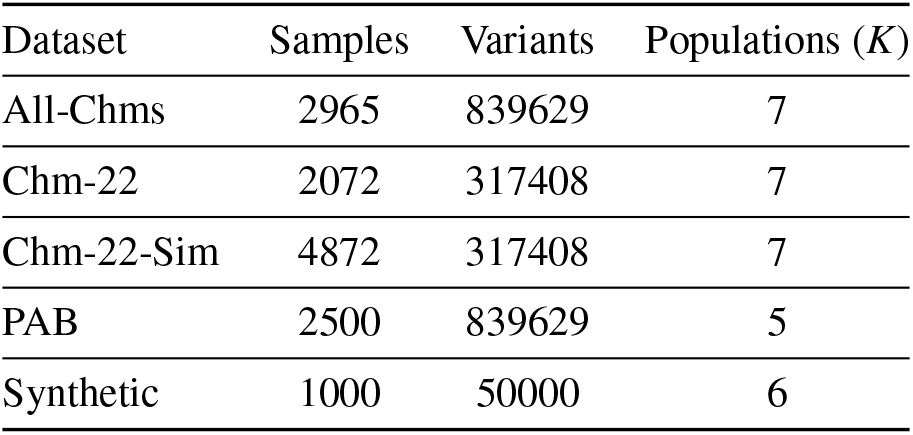
Summary of the datasets used throughout the experiments

**Table 2.** Performance comparison of several global ancestry inference algorithms. Metrics reported from the training data. RMSE(*F*, *F_GT_*) of TeraStructure and HaploNet has not been computed as the allele frequency matrix is not computed in the former algorithm and due to lack of interpretability in the latter. HaploNet was not run on CPU due to resource and time limitations. Runtime format is HH:MM:SS.

| Dataset | Algorithm | $\Delta(Q, Q_{GT})$ | RMSE($Q, Q_{GT}$) | RMSE($F, F_{GT}$) | Runtime (CPU) | Runtime (GPU) |
| --- | --- | --- | --- | --- | --- | --- |
| All-Chms | ADMIXTURE [4, 5] | .042 | .153 | .062 | >1 day | - |
|  | AIStructure [28] | .064 | .159 | .032 | 06:04:28 | - |
|  | TeraStructure [29] | .033 | .133 | - | 02:12:46 | - |
|  | HaploNet [30] | .026 | .114 | - | - | 03:17:00 |
|  | Neural ADMIXTURE | **.025** | **.108** | **.011** | **00:11:21** | **00:01:32** |
| Chm-22 | ADMIXTURE [4, 5] | .048 | .161 | .068 | 02:56:29 | - |
|  | AIStructure [28] | .116 | .256 | .068 | 00:46:49 | - |
|  | TeraStructure [29] | .050 | .170 | - | 00:43:48 | - |
|  | HaploNet [30] | .053 | .170 | - | - | 01:09:29 |
|  | Neural ADMIXTURE | **.033** | **.140** | **.016** | **00:05:46** | **00:00:45** |
| Chm-22-Sim | ADMIXTURE [4, 5] | .046 | .197 | .067 | 09:48:18 | - |
|  | AIStructure [28] | .126 | .286 | .076 | 02:51:36 | - |
|  | TeraStructure [29] | .040 | .175 | - | 06:37:14 | - |
|  | HaploNet [30] | .026 | .113 | - | - | 02:07:54 |
|  | Neural ADMIXTURE | **.011** | **.070** | **6.02e-3** | **00:20:41** | **00:01:34** |
| PAB | ADMIXTURE [4, 5] | **1.44e-4** | **.010** | **5.97e-3** | 03:31:01 | - |
|  | AIStructure [28] | 1.45e-3 | .026 | 7.83e-3 | 05:10:42 | - |
|  | TeraStructure [29] | 1.97e-4 | .012 | - | 01:13:38 | - |
|  | HaploNet [30] | .039 | .248 | - | - | 02:37:09 |
|  | Neural ADMIXTURE | 4.34e-3 | .055 | 7.01e-3 | **00:14:27** | **00:01:48** |
| Synthetic | ADMIXTURE [4, 5] | **1.37e-4** | **.011** | **.028** | 00:08:06 | - |
|  | AIStructure [28] | 2.74e-4 | .014 | .030 | 00:03:07 | - |
|  | TeraStructure [29] | 1.13e-3 | .032 | - | 00:03:28 | - |
|  | HaploNet [30] | .022 | .123 | - | - | 00:04:04 |
|  | Neural ADMIXTURE | 8.60e-4 | .030 | **.028** | **00:01:25** | **00:00:12** |

**Figure 3.**
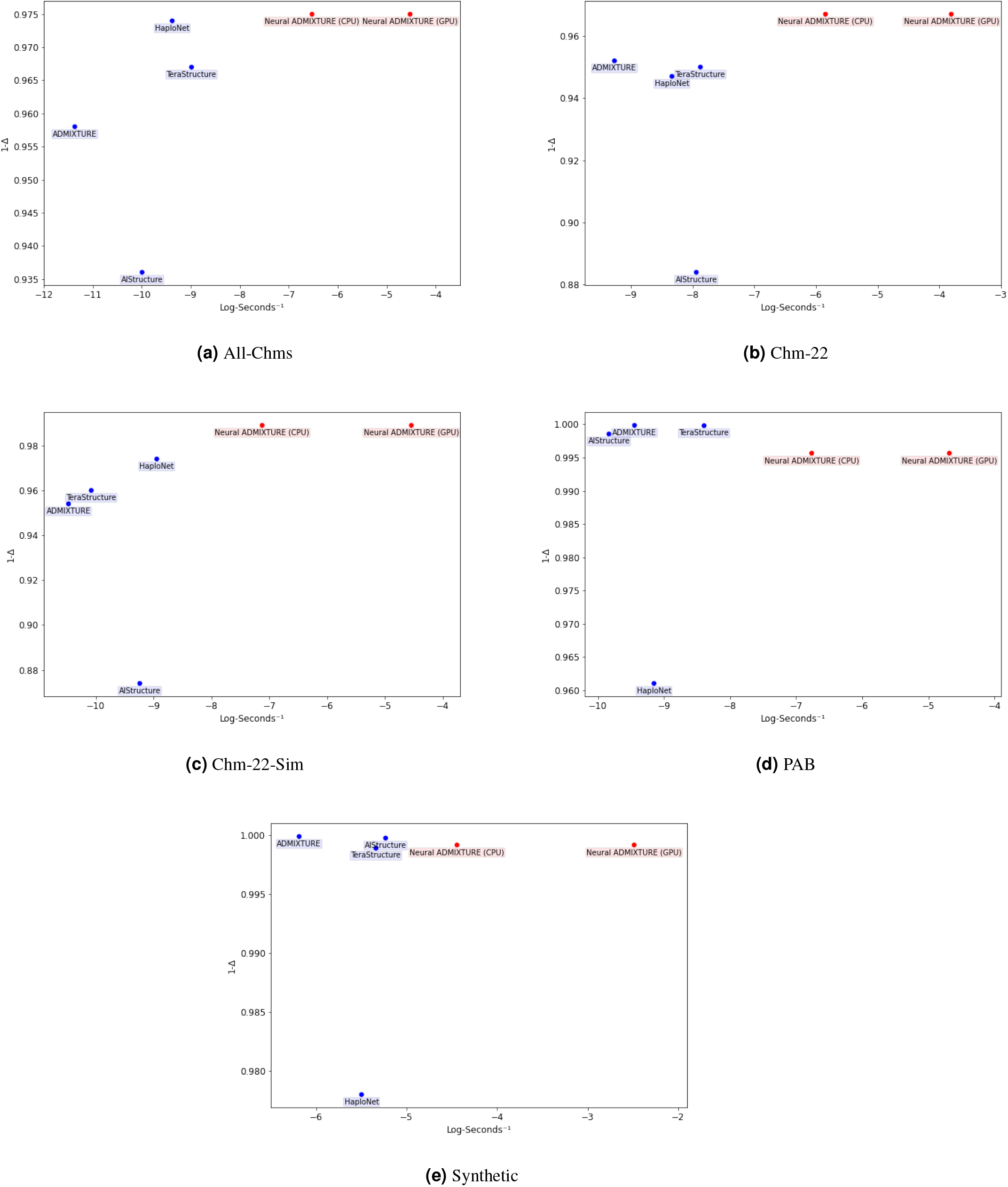
Computation time vs. accuracy plots on five datasets. Note that the X-axis and the Y-axis represent log-seconds^−1^ and 1 – Δ respectively, so faster and better-performing algorithms should be found in the upper-right quadrant. Red is used to highlight our proposed method.

In most datasets, Neural ADMIXTURE also quantitatively outperforms the rest of the algorithms, on both the ancestry assignments themselves (*Q*) as well as the learnt allele frequencies (*F*). In the dataset *All-Chms*, for example, we observe that Neural ADMIXTURE was trained in less than two minutes while ADMIXTURE’s fitting ran for more than a day, with the former surpassing the latter in terms of performance metrics. While Neural ADMIXTURE is not the best-performing algorithm on the datasets PAB and Synthetic, the values of Δ and RMSE in these datasets is very competitive. In Figure 4, it can be observed that on average, Neural ADMIXTURE’s *Q* estimates appear to be more similar to the ground truth matrix than the other *Q* estimations.

**Figure 4.**
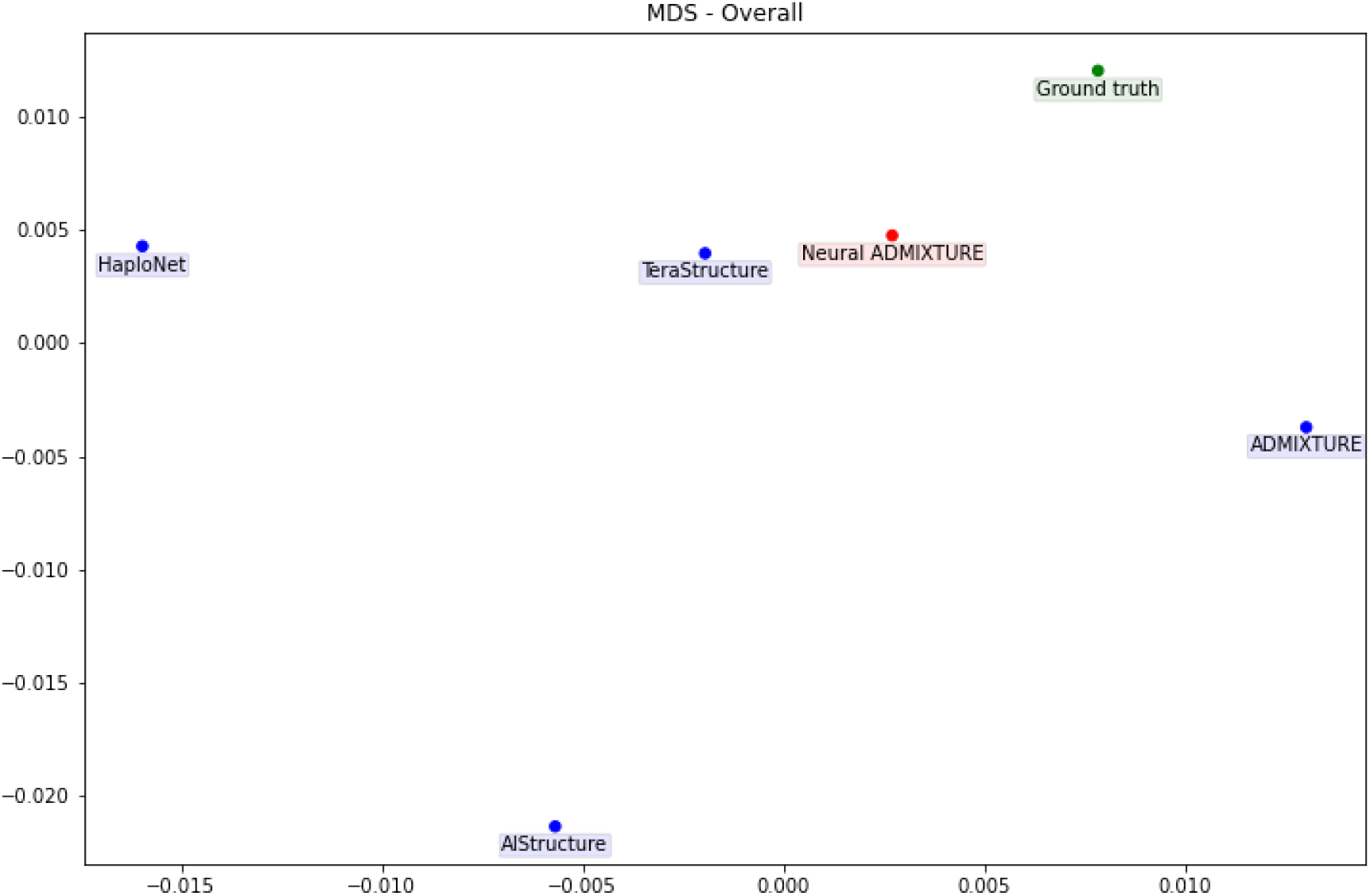
2D visualization of *Q* estimates using multidimensional scaling (MDS). In order to use MDS, a distance matrix between algorithms (including the ground truth matrix) has been computed by using the Frobenius norm between the different *Q* matrices. The average of the distances has been taken among all datasets. Ground truth depicted in green, our proposed method in red.

#### Validation data

Table 3 shows the accuracy and time performance of ADMIXTURE and Neural ADMIXTURE on the validation data of three different datasets. As expected, in the datasets where Neural ADMIXTURE outperformed ADMIXTURE on the training data (*Chm-22, Chm-22-Sim*), Neural ADMIXTURE also outperforms ADMIXTURE on the validation data, and vice versa. Thus, both ADMIXTURE and Neural ADMIXTURE are able to generalize and get meaningful assignments on unseen data.

**Table 3.** Performance comparison of ADMIXTURE and Neural ADMIXTURE on validation data. AlStructue, TeraStructure and HaploNet are not taken into account due to them lacking the possibility of computing ancestry assignments on out-of-training data. Runtime format is HH:MM:SS.

| Dataset | Algorithm | $\Delta(Q, Q_{GT})$ | RMSE($Q, Q_{GT}$) | Runtime (CPU) | Runtime (GPU) |
| --- | --- | --- | --- | --- | --- |
| Chm-22 | ADMIXTURE [4, 5] | .056 | .171 | 00:06:40 | - |
|  | Neural ADMIXTURE | **.043** | **.146** | **00:00:14** | **00:00:07** |
| Chm-22-Sim | ADMIXTURE [4, 5] | .060 | .206 | 00:18:00 | - |
|  | Neural ADMIXTURE | **.025** | **.110** | **00:00:25** | **00:00:07** |
| PAB | ADMIXTURE [4, 5] | **4.29e-3** | **.045** | 00:10:26 | - |
|  | Neural ADMIXTURE | 5.68e-3 | .062 | **00:00:23** | **00:00:10** |

In terms of runtime, Neural ADMIXTURE is again much faster than the classical algorithm on both CPU and GPU, due to the fact that ADMIXTURE must solve the optimization problem again with a fixed *F* to find *Q* for unseen data, while Neural ADMIXTURE directly learns a function Ψ_θ_ (*X*) which estimates *Q*. We note that the inference itself on GPU is extremely fast (generally less than a second to perform a forward pass), but the computational bottleneck in this scenario comes from the reading and processing of the data.

### Qualitative comparison

We also compare ADMIXTURE and Neural ADMIXTURE by visualizing their *Q* estimates and their respective SNP frequencies *F*. Figure 5 contains the prediction over the training and validation data of the dataset Chm-22-Sim. The SNP frequencies (that is, the entries in the *F* matrix) from both models can be observed (projected onto the first two principal components of the training data) in Figure 6. Qualitatively, Neural ADMIXTURE estimates tend to be more polarized, with many samples being assigned only to a single population, while ADMIXTURE appears to be more conservative. On this dataset ADMIXTURE does not differentiate Native Americans (AMR) and East Asians (EAS), and instead partitions Africans (AFR) into two different different ancestry clusters. Neural ADMIXTURE, however, is able to split EAS and AMR populations.

**Figure 5.**
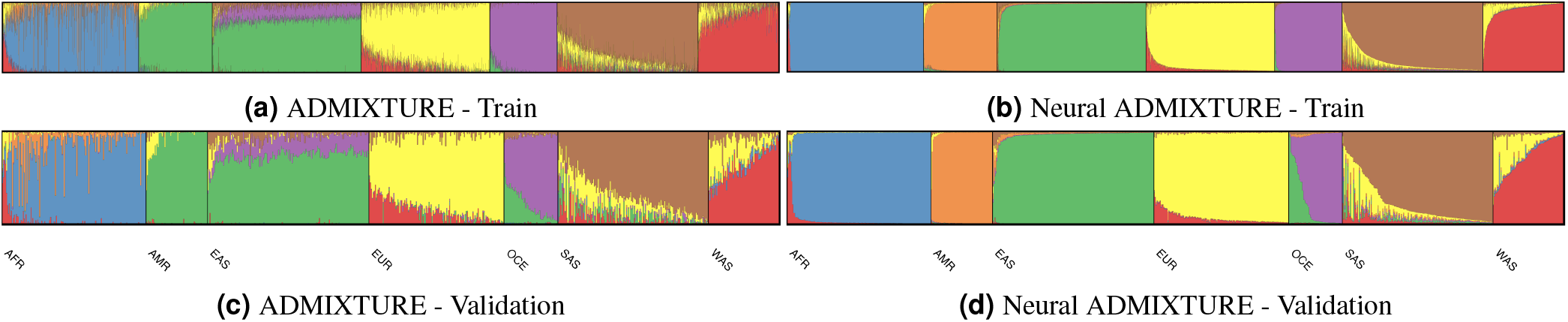
Visualization of *Q* estimates of Chm-22-Sim choosing *K* = 7 clusters. Each vertical bar represents an individual sample and bar color lengths represent the proportion of the sample’s ancestry assigned to that colored cluster.

**Figure 6.**
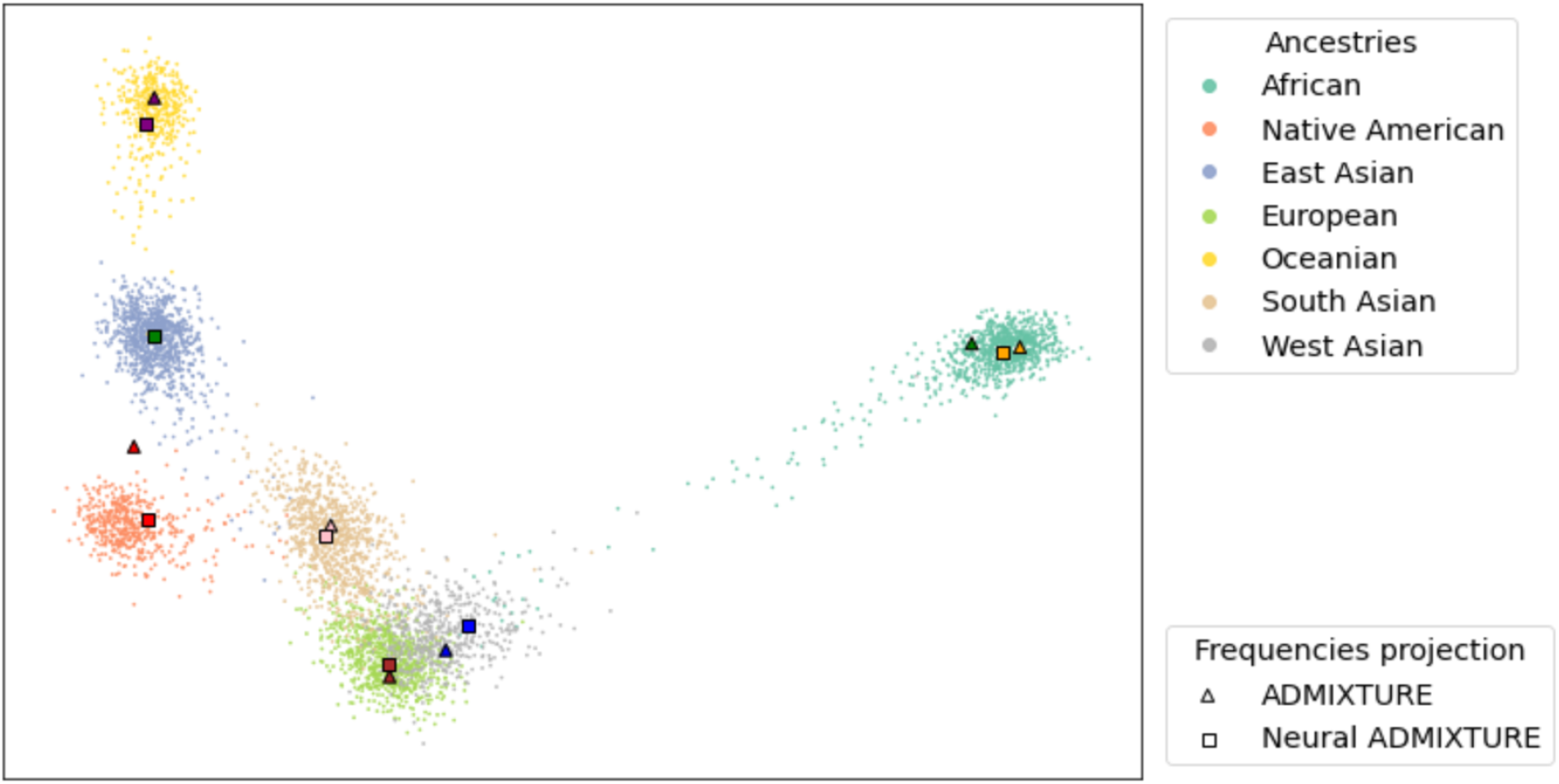
Visualization of *F* matrix of Chm-22-Sim. Training data projected onto its first two principal components along with the projected *F* matrix (cluster centers) from ADMIXTURE and Neural ADMIXTURE.

Such qualitative results are on par with the RMSE and Δ values in both train and validation (Tables 2, 3), indicating that the Neural ADMIXTURE provides estimates which are closer to the ground truth labels as compared to classic ADMIXTURE.

Barplots depicting results of all algorithms on several datasets can be found in the Appendix.

#### Unseen admixed data

In this section, we qualitatively analyze the results of applying Neural ADMIXTURE to unseen admixed individuals: Mexican-Americans (118) and Puerto Ricans (104). To be consistent, we have used the model trained on Chm-22-Sim to retrieve the ancestry assignments.

In Figure 7, we depict results on admixed individuals. Amongst the Mexican-American samples, we observe mainly an orange indigenous component (Native American - AMR) with also a red and yellow component (corresponding to West Asia - WAS and Europe - EUR, respectively). These two latter components likely originate in the immigration of Spanish, Moorish, and Sephardic Jewish individuals during the colonial period. The Puerto Rican samples exhibit EUR, WAS, AFR (African) and AMR ancestry clusters. The additional AFR component corresponds to Afro-Puerto Rican individuals, and is likely linked to the presence of *libertos* during the Spanish conquest of the island, as well as the later introduction of West African slaves to the territory.

**Figure 7.**
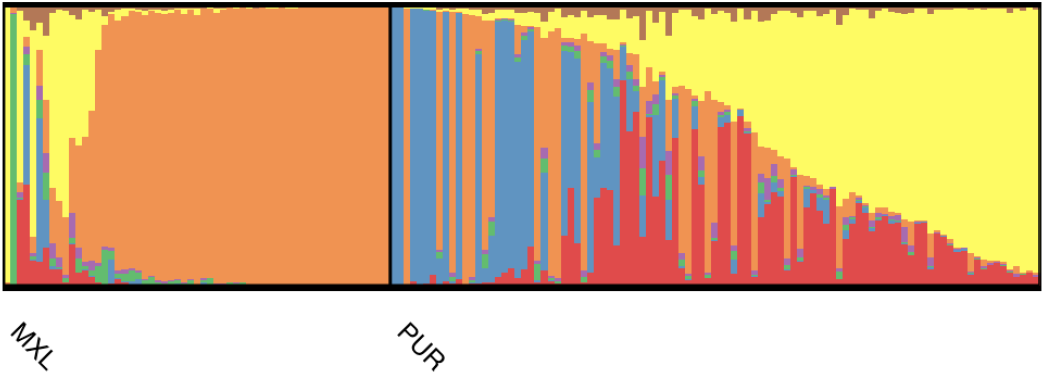
Results on admixed individuals. Individuals of Mexican-American ancestry (MXL, left) and of Puerto Rican ancestry (PUR, right) are shown. The network used to perform inference was trained on Chm-22-Sim.

### Multi-head results

The main advantage of the Multi-head Neural ADMIXTURE (MNA) architecture is that it can perform simultaneous clustering and inference for multiple values of *K* (number of ancestry clusters). Running many different values for *K* allows geneticists to obtain a more complete picture of the variation within populations, and is recommended practice. Figure 8 shows examples of the output of MNA for different clustering results under several values of K ranging from 3 to 9.

**Figure 8.**
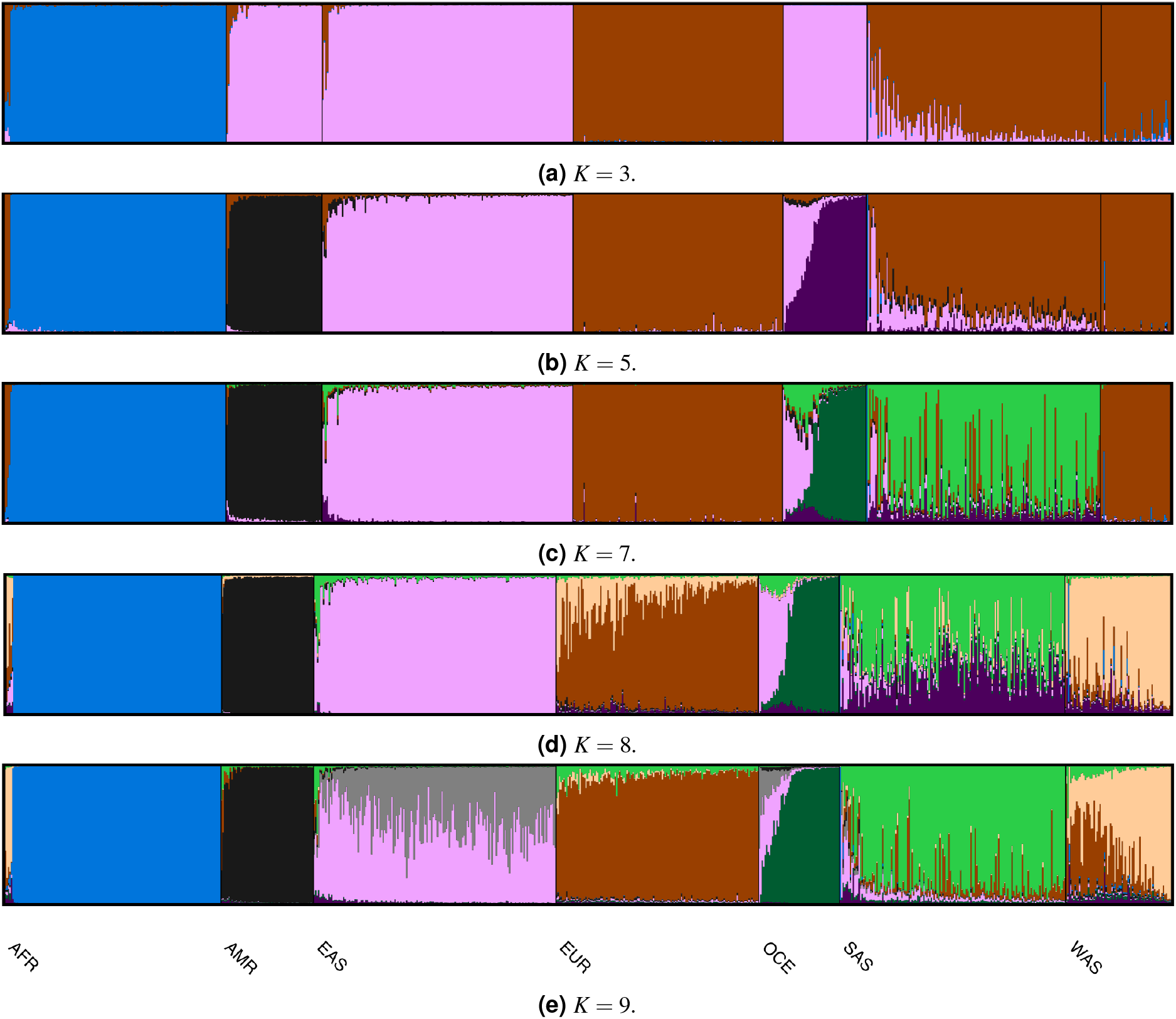
Validation results from Multi-head Neural ADMIXTURE on Chm-22-Sim.

In Figure 8a (*K*=3) we can observe that European (EUR), West Asian (WAS) and South Asian (SAS) are combined within the same cluster, while Oceanian (OCE) and East Asian (EAS) are clustered together, and African (AFR) has its own cluster. These results reflect the genetic similarity between the respective groups due to their Out-of-Africa migration patterns and subsequent gene flow.

After increasing to *K* = 5, OCE obtains its own cluster, reflecting the ancient divergence of that population from the others. As more clusters are incorporated, Native American (AMR) and EAS obtain their own clusters and OCE is divided between a component found predominantly in OCE and a component characteristic of EAS. The latter likely reflects the later migration of Austronesian speakers from East Asia out into the Pacific Islands, where they contributed their ancestry to the Oceanian inhabitants. A shared component between EUR, SAS and WAS is maintained, independent of the cluster number *K*, which could be linked to early farmer expansions out of West Asia and into both Europe and South Asia, following the birth of agriculture, as well as to the much later expansion of the Indo-European language family across all of these regions. Other genetic exchanges between these neighboring regions doubtlessly played a role.

With a sufficiently high number of clusters (Figure 8e) a shared component between WAS and some AFR populations appear, which might reflect North African gene flow.

## Scalability analysis

In order to assess the scalability of different methods, we simulated multiple datasets with several number of variants and number of samples using the software provided in [27]. The datasets consist of all combinations of *N* ∈ {1000, 5000, 10000, 20000, 50000} and *M* ∈ {1000, 10000, 50000, 100000}, where *N* is the number of samples and *M* is the number of SNPs.

We compare the training times of ADMIXTURE, AlStructure, TeraStructure and our proposed method, Neural ADMIXTURE (both on CPU and GPU). Figures 9a and 9b depict the evolution of the execution times when fixing the number of samples (10000) and SNPs (100000) respectively. Plots for other values of *N* and *M* can be found in the Appendix, as well as the hyperparameters used.

**Figure 9.**
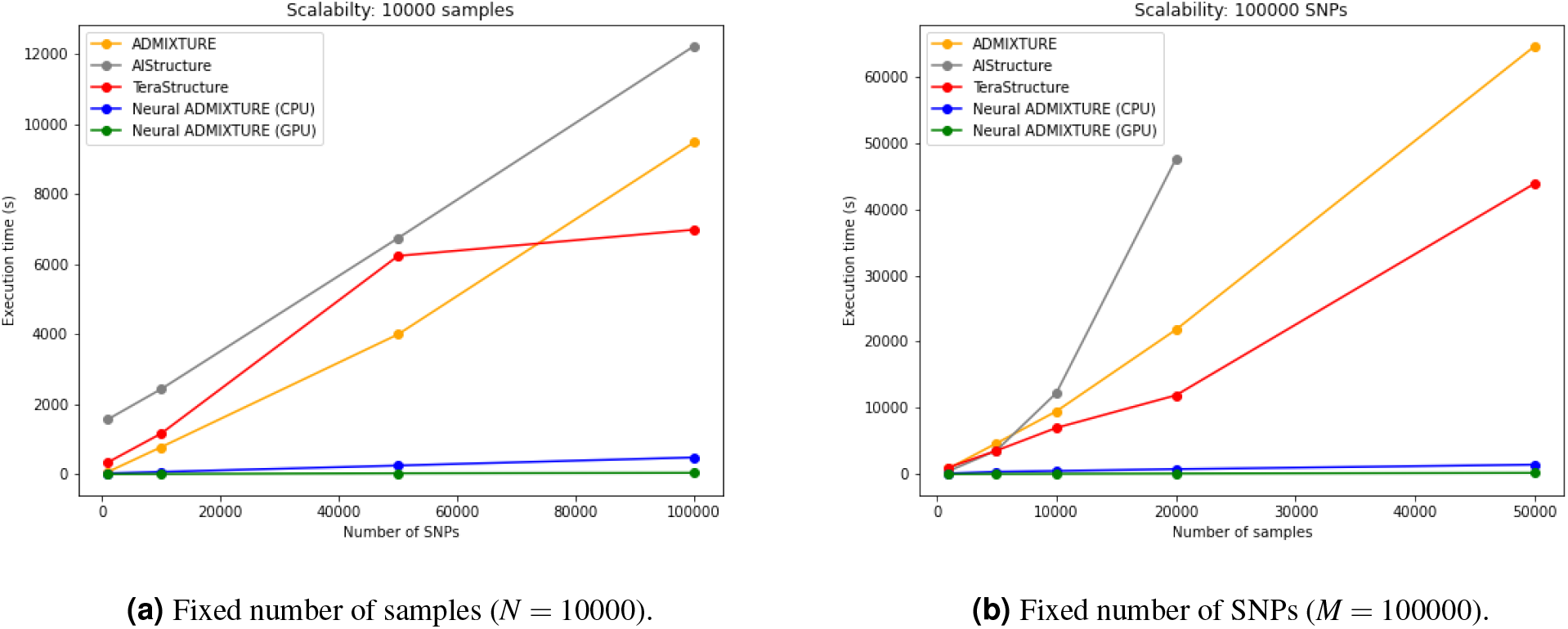
Evolution of execution time when increasing number of samples and number of variants. AlStructure results are not reported on 50000 samples due to exceedingly high execution time.

Results seen in Figures 9a and 9b show that Neural ADMIXTURE is consistently much faster than ADMIXTURE, AlStructure and TeraStructure, especially for larger number of samples, which is the area of application of the latter two as well as Neural ADMIXTURE. Moreover, and not surprisingly, Neural ADMIXTURE scales even better than the other methods using GPUs.

## Discussion

In this paper, we demonstrate that the unsupervised clustering algorithm ADMIXTURE can be re-framed as an autoencoder. This novel framing, which we name Neural ADMIXTURE, allows for the use of common neural network optimization techniques such as SGD or Adam and provides rapid inference through the encoder, two orders of magnitude faster than the original ADMIXTURE algorithm as well as other similar algorithms. Furthermore, by adding more heads, multiple estimates with different priors on the cluster number (*K*) can be performed simultaneously, reducing overall compute time still further. This approach, combined with the use of GPU compute, can enable rapid and accurate results on even large modern biobanks.

Neural ADMIXTURE relates heavily to amortized inference, in which already computed inferences are reused to calculate inferences on novel data [32] by trying to learn a function which computes the parameters rather than the parameters themselves. While more classical amortized inference techniques may be explored in order to improve execution times of some algorithms, we believe that the speedup will not be as massive as the one introduced by Neural ADMIXTURE, mainly due to the acceleration provided by the neural architecture themselves as well as the usage of GPUs.

Taking into account the fact that several runs are typically required to find optimal settings, the multi-head architecture as well as the naturally fast execution time translates into being able to run many executions of the software with different hyperparameters, making this optimality search much less time-consuming. Moreover, the available software provides tools to easily run hyperparameter sweeps and visualize the results, making the process even less tedious.

Finally, we note that the two proposed initializations: PCK-Means (Algorithm 1) and PCArchetypal (Algorithm 2) work best in different scenarios. Due to the nature of K-Means and Archetypal Analysis, if the data consists mainly of single-ancestry individuals, PCK-Means tends to give better results that PCArchetypal, and vice-versa if the data contains mainly admixed individuals. Inspecting a two-dimensional projection of the SNP data (also integrated in the software) usually suffices in order to decide which initialization might work better. Nevertheless, the low computational complexity of our proposed method should allow several runs with both initialization algorithms.

## Acknowledgments

This work was supported in part by NIH grant 7U01HG009080, and project PID2020-117142GB-I00, funded by MCIN/ AEI /10.13039/501100011033.

## Appendix A: Training results

This section contains bar plots depicting training results on most datasets described in the Experiments section. Visualizations have been obtained using the software *pong* [33].

### All-Chms

**Figure 10.**
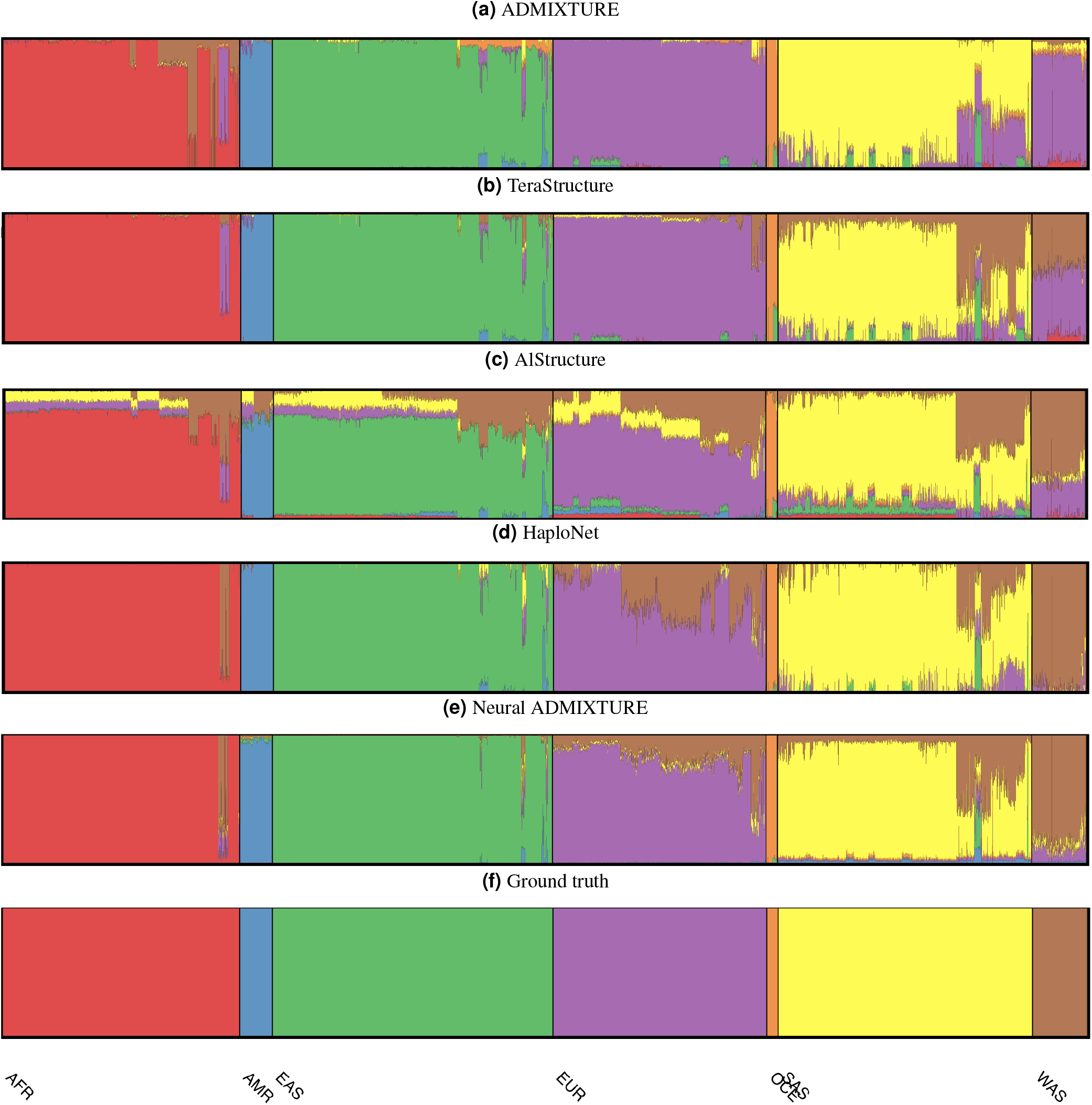
*Q* estimates on All-Chms.

### Chm-22

**Figure 11.**
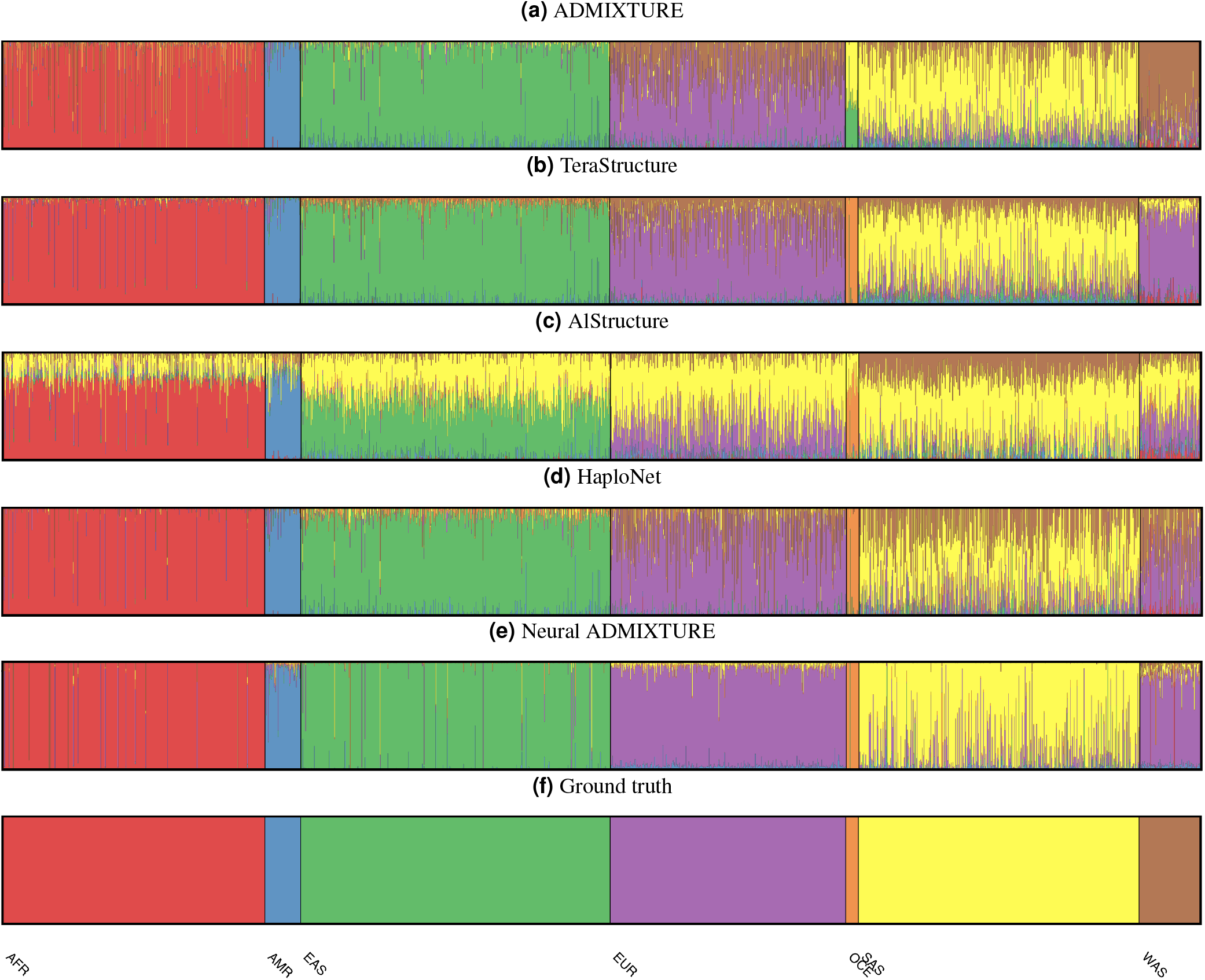
*Q* estimates on Chm-22.

### Chm-22-Sim

**Figure 12.**
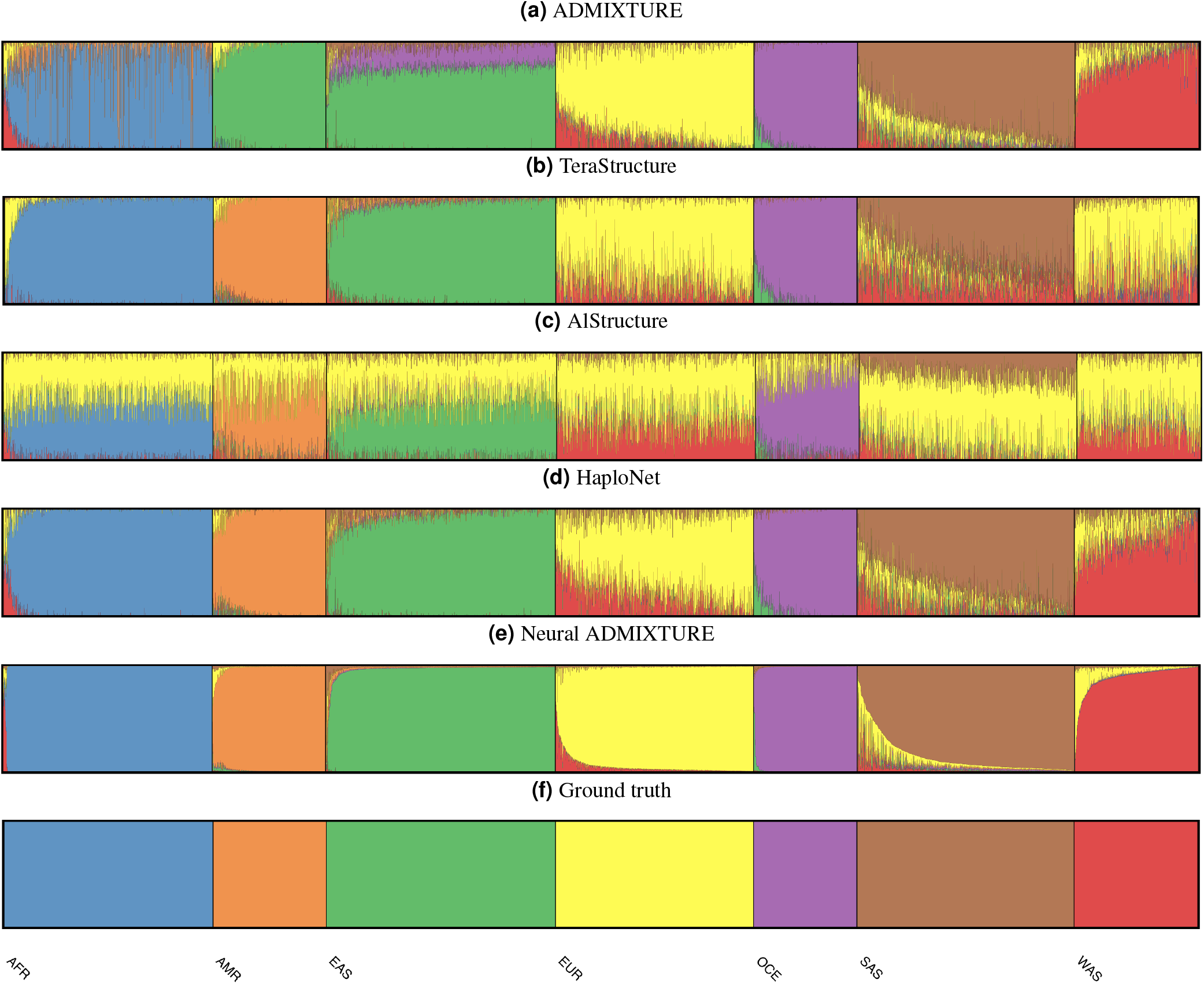
*Q* estimates on Chm-22-Sim.

### PAB

**Figure 13.**
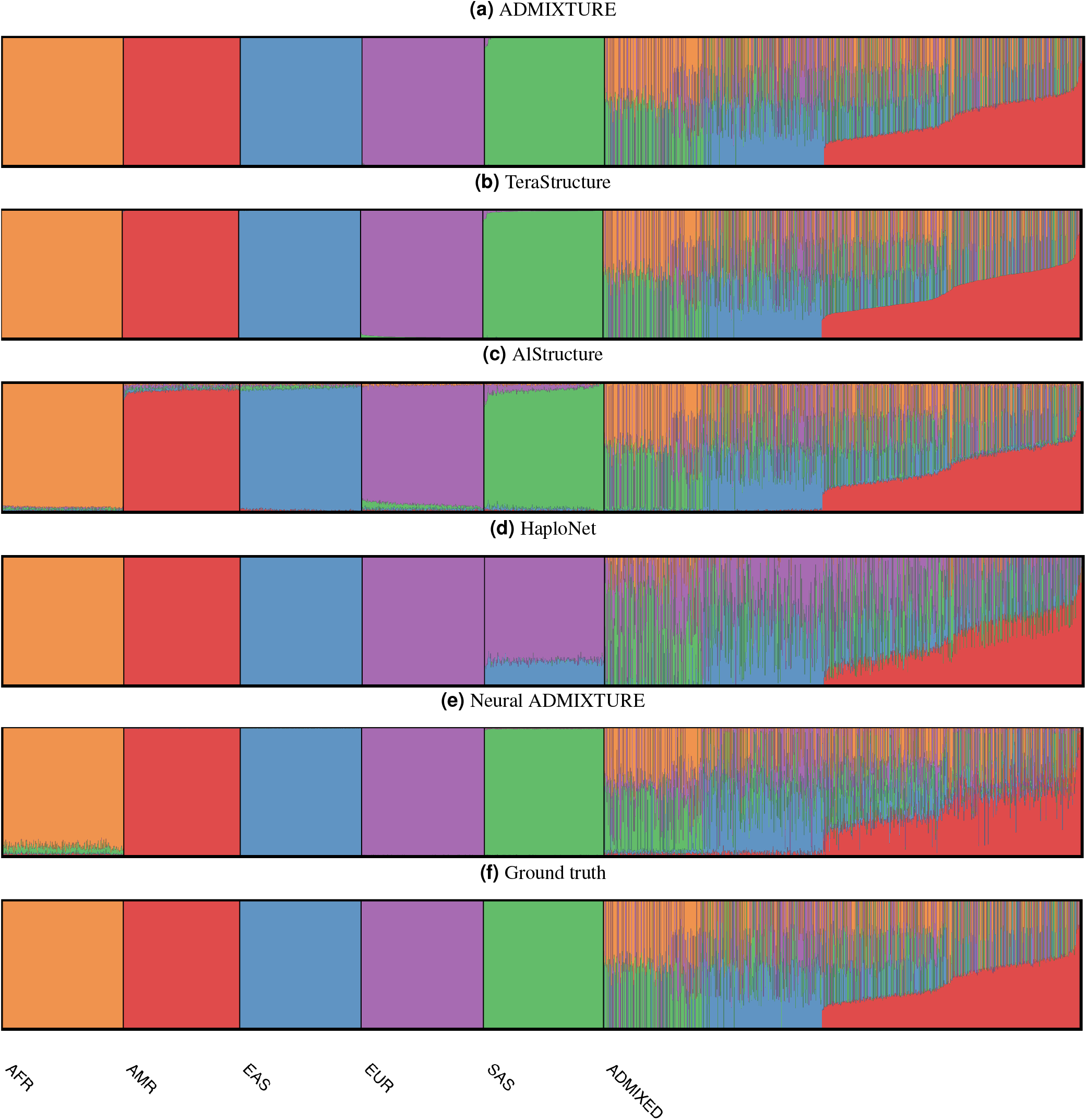
*Q* estimates on PAB.

## Appendix B: Hyperparameters

All algorithms have been run with the hyperparameters suggested by the respective authors.

ADMIXTURE and AlStructure do not have critical hyperparameters to tune, and the defaults of the implementation are used.

In the case of TeraStructure, the most critical parameter is the frequency of convergence checks, which has been set to 7% the number of variants (authors suggest using some value between 5%-10%).

HaploNet has been trained using a batch size of 4096, as well as a window size of 128 for 30 epochs per window.

For Neural ADMIXTURE, a batch size of 400 is used, as well as an L2 penalty (λ) of 5e-4. Eight principal components have been used in the initialization step on all datasets. The other hyperparameters can be found in Table 4.

**Table 4.** Hyperparameters used to train Neural ADMIXTURE on different datasets.

| Dataset | Initialization | Epochs | Learning rate |
| --- | --- | --- | --- |
| All-Chms | PCK-Means | 3 | $1e-5$ |
| Chm-22 | PCK-Means | 7 | $1e-4$ |
| Chm-22-Sim | PCK-Means | 13 | $1e-4$ |
| PAB | PCArchetypal | 5 | $1e-5$ |
| Synthetic | PCArchetypal | 24 | $1e-4$ |

## Appendix C: Scalability analysis

In this section we provide the results of the scalability analysis mentioned in the Experiments of the main text.

The hyperparameters used to fit ADMIXTURE, AlStructure and TeraStructure are the same as described in Appendix B. Neural ADMIXTURE has been trained using a learning rate of 1*e* – 4 and the PCArchetypal (Algorithm 2) initialization.

**Figure 14.**
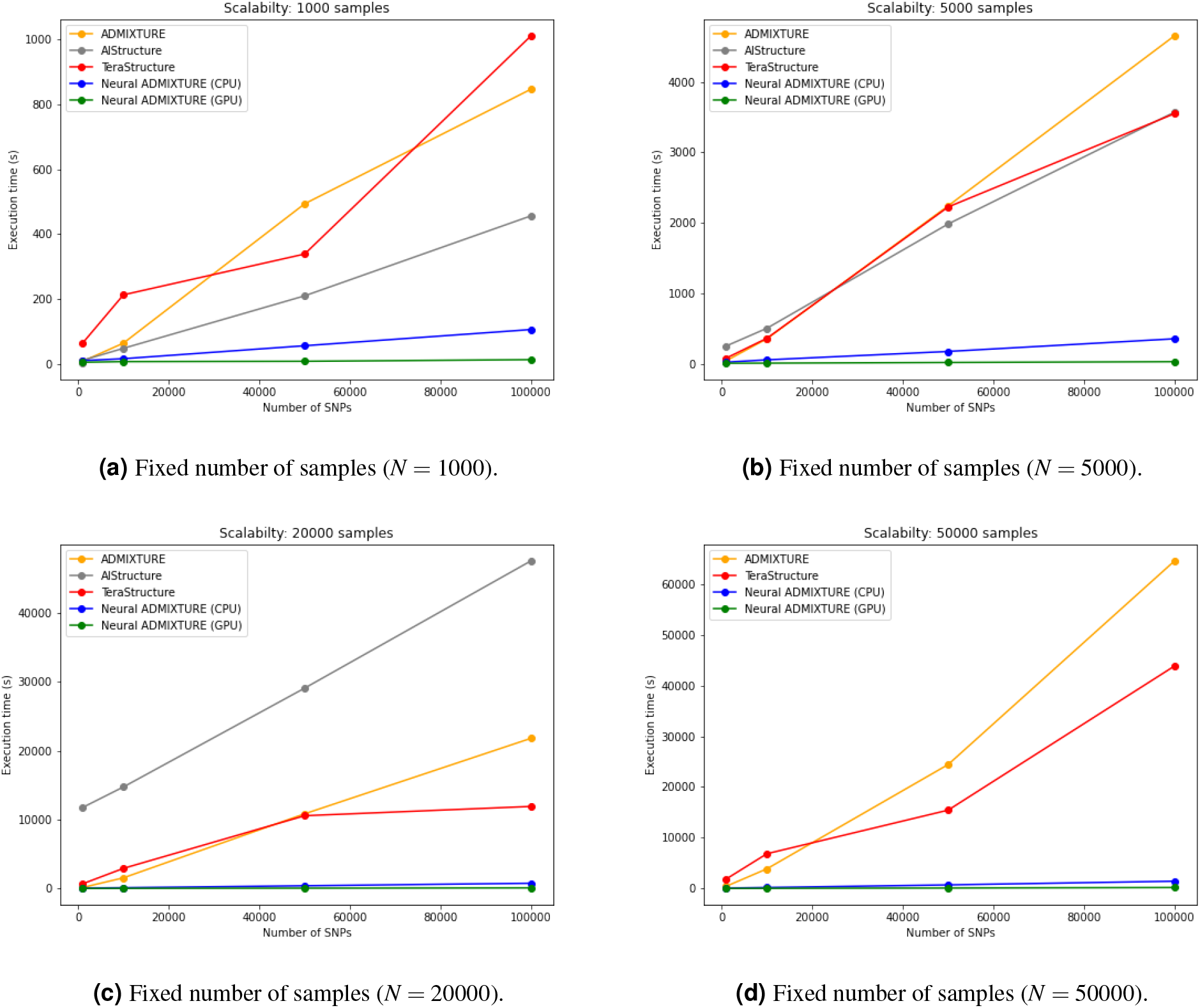
Evolution of execution time when increasing number of samples. AlStructure results are not reported on 50000 samples due to exceedingly high execution time.

**Figure 15.**
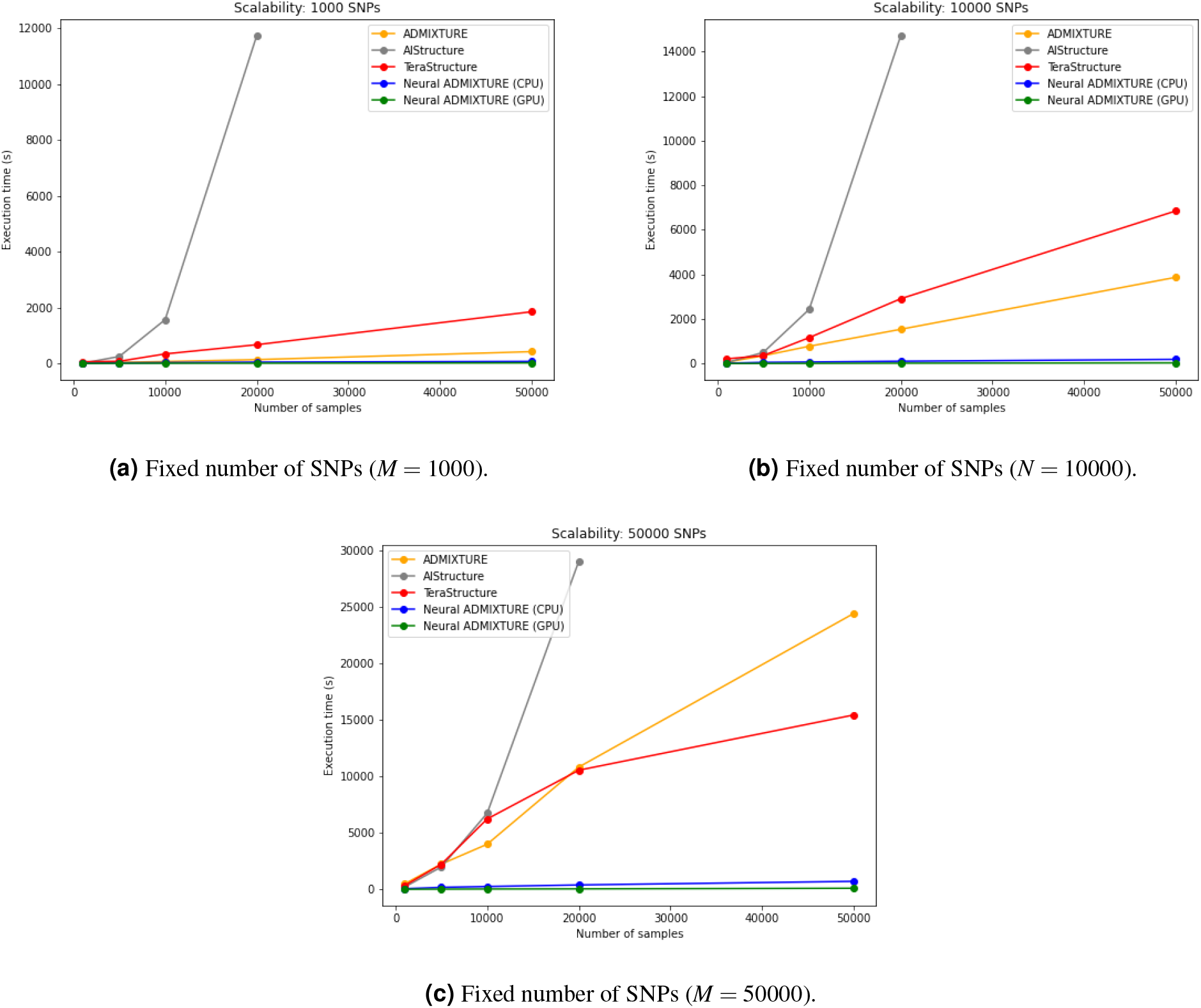
Evolution of execution time when increasing number of SNPs. AlStructure results are not reported on 50000 samples due to exceedingly high execution time.

